# Identification and Characterization of Interacting Proteins of TARANI/Ubiquitin Specific Protease-14 in *Arabidopsis thaliana*

**DOI:** 10.1101/2025.02.01.636027

**Authors:** Anjana S Hegde, Utpal Nath

## Abstract

Ubiquitin proteases play a crucial role in protein degradation and turnover by regulating the cleavage of polyubiquitin chains. TARANI/UBIQUITIN SPECIFIC PROTEASE-14 (TNI/UBP14) specifically cleaves Lys-48-linked and linear polyubiquitin chains into mono-ubiquitins. The *tni* mutant exhibits pleiotropic phenotypes, including cup-shaped leaves, tri-cotyledons, reduced lateral roots, and increased petal number, though the underlying mechanisms driving these phenotypes remain unclear. In this study, we generated TNI transgenic lines and employed immunoprecipitation mass spectrometry, proximity labelling, and yeast two-hybrid screening to identify TNI’s interacting proteins. These analyses revealed 92 interactors involved in diverse biological processes, including protein and carbohydrate metabolism, light signalling, and intracellular transport. Subcellular localization analysis showed that many of the interacting proteins are located in the nucleus and cytoplasm, suggesting that TNI’s nuclear localization may regulate gene function. We further validated the *in planta* biological significance of ULTRAPETALA 2 and HASPIN KINASE as key interacting partners of TNI. These findings uncover previously uncharacterized functions of TNI/UBP14, shedding light on its central role in cellular processes and providing insights into its regulatory mechanisms—an area that has remained largely unexplored until now.

**Summary statement:** The proteins that interact with the TARANI/ Ubiquitin protease 14 in vivo have been identified using immunoprecipitation mass-spectrometry methods. Identification of non-overlapping targets highlight the importance of using diverse protein identification methods.

## Introduction

Proteins, the building blocks of the cell, are abundant in the cell. They often function in complex with other proteins. Therefore, it is essential to maintain protein homeostasis through the continuous recycling process. The specialized 26S proteasomal machinery ensures cellular protein homeostasis through continuous protein turnover, involving both ubiquitylation and deubiquitylation processes. Ubiquitylation, a crucial protein modification, governs various cellular processes. The coordinated action of E1 activating enzyme, E2 conjugating enzyme, and E3 ubiquitin ligase tags target proteins for degradation by adding ubiquitin. De-ubiquitylation reverses the E3 ligase function by hydrolyzing ubiquitin bonds(Clague, 2019).

The ribosomal protein-anchored single ubiquitin or poly pro-ubiquitin chain is the de novo synthesis mode of ubiquitin. Iso-linked poly-ubiquitins are produced by cleaving the poly-ubiquitin chains from the target proteins during proteasome-mediated degradation. Deubiquitinases (DUBs) hydrolyse poly-ubiquitins and cleave ribo-anchored mono-ubiquitins to replenish the mono-ubiquitin pool(Komander et al., 2009). Arabidopsis genome encodes 64 DUBs, of which 27 are ubiquitin-specific proteases (UBPs) (Zhou et al., 2017). UBPs are specialised in function to hydrolyze ubiquitins from the peptide or iso-peptide bonds (Sullivan et al., 1990). Single mutants of 25 out of 27 members showed no visible phenotypes suggesting functional redundancy. A transfer DNA (T-DNA) mediated single mutants of *ubp14* and *ubp19* were embryonic lethal (Liu et al., 2008; Doelling et al., 2001). *ubp15* mutant exhibited a defect in cell proliferation, resulting in narrow and serrated leaves (Liu et al., 2008). Higher order double knockout *ubp12-1 ubp13-3* exhibited severe phenotypes like small plants, round leaves, dwarfism, and more branches after bolting compared to individual parents (Cui et al., 2013).

Mutants of the proteasomal system proteins showed pleiotropic effects in plants. An amino acid substitution in the RPT5a subunit of the 26S proteasome led to abscisic acid and ethylene-sensitive, and light hyposensitive mutants (Hayashi & Hirayama, 2016). The *rpn1a* mutation resulted in embryonic lethality in homozygous lines displaying arrest at the globular stage with defects in the embryonic root, protoderm, and procambium (Brukhin et al., 2005).

A forward genetic screen aimed at identifying mutants with altered leaf morphology in Arabidopsis identified the *tarani* (*tni*) ["boat" in Sanskrit] mutant, which exhibits a cup-shaped leaf phenotype (Karidas et al., 2015). *TNI* codes for ubiquitin-specific protease 14 (UBP14) which cleaves α-linked and isopeptide-linked poly-ubiquitin chains into mono-ubiquitin (Majumdar and Karidas et al., 2020). Arabidopsis genome contains a single copy of *UBP14*, and the null alleles of *UBP14* are embryonic lethal, suggesting its evolutionarily conserved role(Tzafrir et al., 2002; Majumdar & Nath, 2020). G to A mutation in the intron 3 of the *tni* mutant led to the retention of the intron 3 in nearly 50% of mRNA transcripts due to reduced enzymatic efficiency of the spliceosomes. However, it did not affect the protein synthesis in the *tni* mutant, but the *tni* mutant possibly consists of both mutant and wild-type proteins (Majumdar and Karidas et al., 2020). Deletion of UBP14 in *Saccharomyces cerevisiae* and *Dictyostelium discoideum* showed that UBP14 homologs are not essential for the survival of these organisms (Doelling et al., 2001; Lindsey et al., 1998).

AtUBP14 has an N-terminal ZnF-UBP domain followed by a protease domain and two consensus C-terminal Ubiquitin Associated (UBA) domains (Majumdar & Karidas et al., 2020). The ZnF-UBP and UBA domains serve as interfaces in protein-protein interaction (Ravasi et al., 2003). UBP14 has been shown to localize to the nucleus and regulate endoreduplication by forming a complex with the nuclear-localized ULTRAVIOLET INSENSITIVE-4 (UVI4) (Xu et al., 2016). In *Magnaporthe oryzae* UBP14-GFP pulldown, an Arabidopsis UBP14 homologue showed interaction with multiple proteins which are involved in functions like carbohydrate metabolism, stress response, membrane transportation, secondary metabolism, and ubiquitin system (Wang et al., 2018). The compromised proteasomal activity in the *tni* mutant increases AUX-IAA transcriptional repressors’ stability, leading to reduced auxin response (Majumdar & Karidas et al., 2020). The ZnF-UBP domain in TNI led us to hypothesize that UBP14, central to the ubiquitin-proteasome system (UPS), probably interacts with multiple proteins to regulate various cellular processes, resulting in the pleiotropic phenotypes of its mutants.

Ubiquitin proteasomal components play multiple roles. They also regulate various cellular activities besides their function in protein degradation. These include transcriptional regulation, epigenetic control, flowering, immune response, tissue growth, stress adaptation, and nutrient homeostasis within the cell (An et al., 2018; Cho et al., 2006; Cui et al., 2013; Ewan et al., 2011; Ezhkova & Tansey, 2004; Ferdous et al., 2002). UBP members interacted with proteins involved in various biological functions. UBP13 stabilises ROOT MERISTEM GROWTH FACTOR 1 (RGF1)-PERCEIVING RECEPTOR (RGFR1) by deubiquitination and regulates the transcription factors PLETHORA1/2(PLT1/2) in root meristem development in Arabidopsis (An et al., 2018). UBP16 interacts with SERINE HYDROXYMETHYLTRANSFERASE1 (SHM1) and SHM4 *in vitro* and *in vivo* to regulate salt stress and stabilisation of SHM1 by deubiquitination which confers salt tolerance by maintaining the proper ratio of salts in the cells through Na+/K+ transporters (Zhou et al., 2013). UBPs have also been shown to mediate epigenetic modification in partnership with other polycomb group proteins (PcG). UBP12/13 associates with LIKE HETEROCHROMATIN PROTEIN 1 (LHP1), a plant-specific polycomb protein, *in vivo* and is required for H3K27me3 to repress the polycomb target genes FLOWERING LOCUS C (FLC), MADS AFFECTING FLOWERING4 (MAF4) and MAF5. UBP12 deubiquitinates H2A, and H2A ubiquitination brings out gene silencing of some PcG proteins (Derkacheva et al., 2016). This suggests that UBP 12/13 are involved in the balancing act of epigenetic regulation of flowering.

Protein-protein interactions (PPI) are fundamental for cellular function. Over the past two decades, proteomic studies have evolved from whole-cell analyses to organelle-specific, tissue-specific and single-cell proteomics.

The advancement in protein tagging techniques enabled the identification of organelle and tissue-specific low-expressed proteins. To tackle the challenges associated with studying the stochastic interactions in the biological system, a modified version of the biotin ligase BioID from *E.coli* was developed to map protein interactions (Roux et al., 2012). The feasibility and non-lethal nature of biotin led to its rapid popularity, making it one of the most preferred techniques for studying protein interactions, even in other biological systems like mammalian and plant cells (Kim et al., 2016). TurboID, the modified version of BioID, is efficient in labelling yeast, flies, worms, chickens, mammals, and plant cells within 10 minutes (Branon et al., 2018). The task of identifying nuclear membrane interactors was made easy by BioID tagging (Huang et al., 2020). Proximity labelling made these studies possible with minimal tissue and without much sample loss (Roux et al., 2012; Kim et al., 2016; Branon et al., 2018). Since cellular processes are dynamic, using more than one method to study the protein-protein interactions is preferable. The constant turnover or the dynamic activity of the proteins in the cell results in variation in the immediate environment.

In this study, we set out to uncover the comprehensive interactome of TNI by employing multiple approaches. We identified the interactors of TNI by immunoprecipitation-mass spectrometry (IP-MS), proximity labelling, and yeast two-hybrid screening. We validated a selected subset of interactions from our high throughput screen by binary methods and conducted a focused investigation on the biochemical and genetic implications of these interactions using ULTRPETALA 2 and HASPIN KINASE as a case study.

## Results

### Generation of TNI transgenic lines

We generated transgenic Arabidopsis lines expressing *TNI* with a C-terminal Myc tag under CaMV *35S* promoter, *RPS5A* promoter, and endogenous *TNI* promoter in the Col-0 background. A detailed phenotypic analysis of these transgenic lines in the T2 generation revealed that nearly 20% of the transgenic plants show cup-shaped leaves similar to the *tni* mutant (Fig. 1A). Further, expression of the transgenic TNI protein was confirmed using western blot analysis (Supp Fig. 1A). TNI protein expression was also visualized in different tissues of *pTNI::TNI-GUS* transgenic line. GUS assay showed that TNI is expressed almost throughout the plant and at all stages of plant growth, as was reported earlier (Doelling et al., 2001).TNI-GUS signal was detected throughout the plant in the seedling stage (Fig. 1B). Since protein expression was observed throughout the seedling, we chose 8-day-old seedlings for the *in vivo* protein interaction identification experiments. TNI was previously shown to be localised to the nucleus in the Nicotiana leaves when over-expressed under *35S* promoter (Xu et al., 2016). We generated a transgenic line expressing the TNI-G3GFP protein to check the subcellular localisation of TNI in the cell. TNI was found to be localised to both the nucleus and cytoplasm (Fig. 1C). Previously, it was reported that the lateral root number is significantly less, and hypocotyl is two times longer in the *tni* mutant than in the Col-0 (Majumdar and Karidas et al., 2020). The lateral root number was significantly higher, and hypocotyl length was significantly longer in the TNI overexpression transgenic lines (Fig. 1D, E). The belowground phenotypes of TNI transgenic lines contrast with that of the *tni* mutant. With this phenotypic and biochemical analysis confirmed, we confirmed the TNI transgenic lines for further *in vivo* studies.

**Figure 1.**
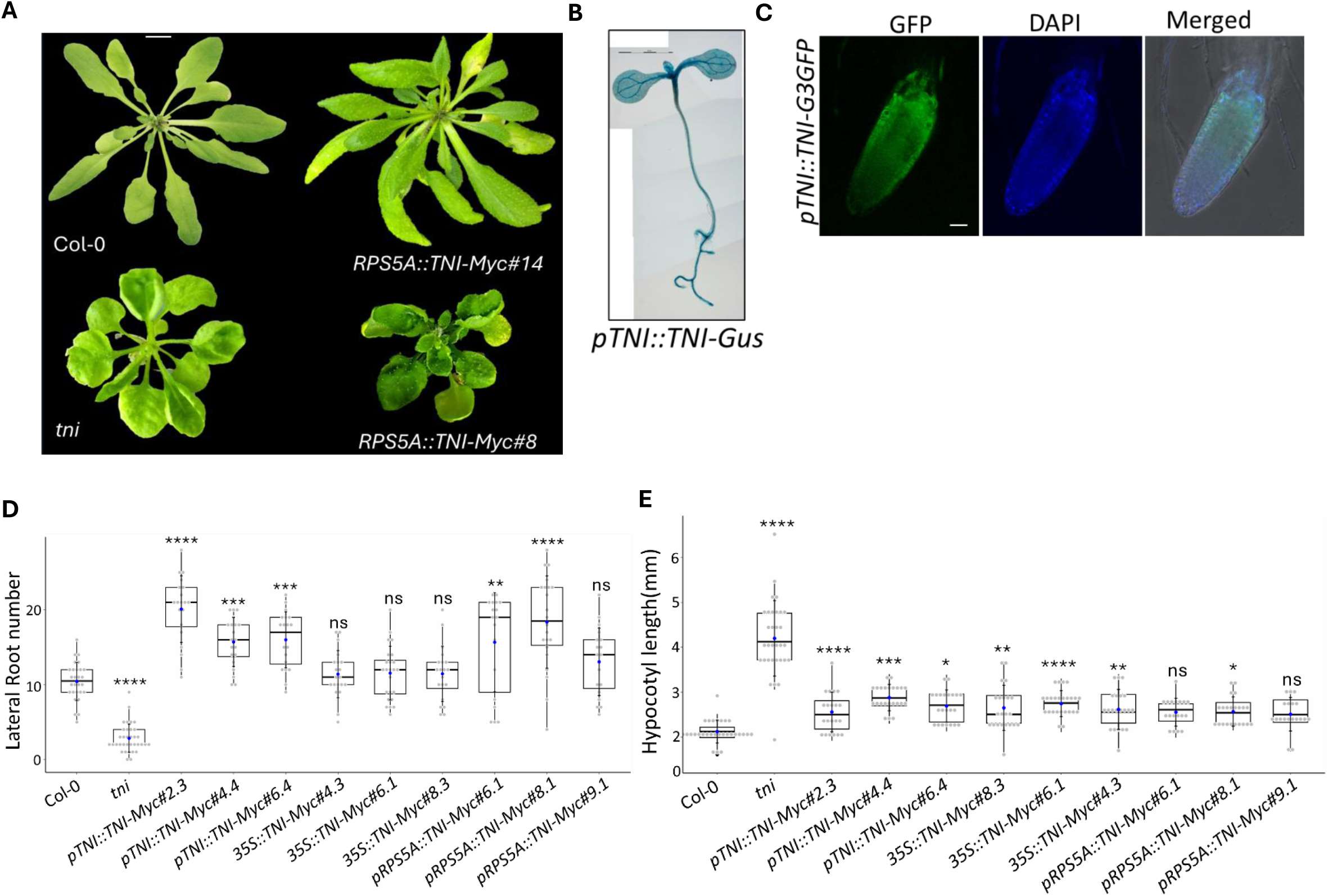
Phenotypic evaluation of *TNI* transgenic lines. (A) Top view of 30-day old rosettes of the indicated genotypes. Two independent transgenic plants overexpressing TNI-Myc fusion protein (*RPS5A::TNI-Myc#8 & RPS5A::TNI-Myc#14* are shown. Scale bar, 1 cm. (B) An 8-day old *pTNI::TNI-GUS* seedling processed for GUS activity. (C) Root tips of 8-day old *pTNI::TNI-G3GFP* seedlings showing TNI-GFP signal. DAPI staining visualises nuclei. (D-E) Average number (n = 25-30) of lateral roots (D) and average length of hypocotyl (E) in 12-day old (D) and 8-day old seedlings, respectively in the indicated genotypes. (). Error bars represent SD. Asterisks indicate significance using Tukey’s *t*-test (**** *p*<0.0001, \*\*\**p*<0.001, \*\**p*<0.01, \**p*<0.1, ns, not significant). Data are shown for nine independent transgenic lines.

### Identification of interacting proteins of TNI

#### a. Immunoprecipitation-MS (IP-MS)

We selected the *RPS5a::TNI-Myc #6* transgenic line to perform an immunoprecipitation-pulldown experiment. The total protein extract from the *RPS5a::TNI-Myc* seedlings was pulled down against anti-Myc and anti-IgG antibodies (Fig.2A). We conducted experiment with three biological replicates, including respective controls, and confirmed the equal amount of TNI in all the replicates (Supp Fig. 1B). The presence of TNI was validated by anti-Myc immunoblot in both the anti-Myc and anti-IgG pulldown (∼120kDa) (Fig. 2B). We found 29 proteins enriched in the anti-Myc pull-down against the IgG negative control (Fig. 2C and Table 1).

**Figure 2.**
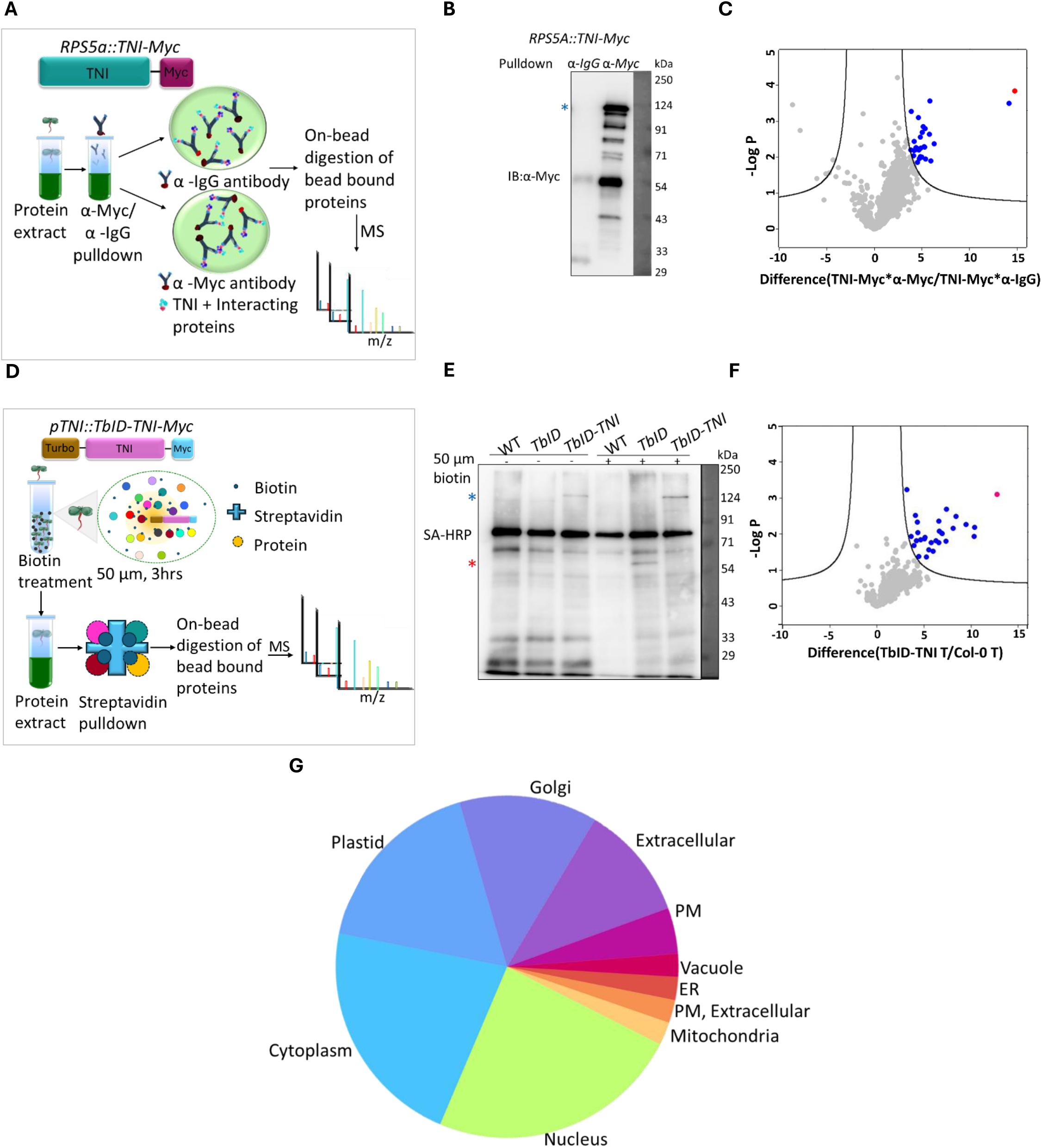
Identification of TNI-interacting proteins by immunoprecipitation mass spectrometry (IP-MS) and affinity purification (AP-MS). (A) A schematic description of the IP-MS experimental setup. Immuno-pulldown was performed by incubating total protein extract from *RPS5A::TNI-Myc* tissue with α-Myc and α-IgG (negative control) antibodies, followed by MS analysis. (B) A representative western blot image of TNI-Myc bait protein after pulldown with α-IgG or α-Myc antibodies. Experiment was repeated three times and similar results obtained. Blue asterisk corresponds to the TNI-Myc band. Numbers on the right indicate molecular weight. (C) Volcano plot showing proteins enriched in α-Myc pulldown sample over the α-IgG control in the IP-MS experiment described in (A-B). Blue circles, enriched proteins; red circle, TNI-Myc protein. (D) A schematic representation of the AP-MS experimental setup. Total protein extracts from 8-day old seedlings expressing *pTNI::TbID-TNI-Myc* and *pTNI::TbID-Myc* (negative control, not shown here) were treated with biotin, pulled-down by pulldown using streptavidin beads, followed by MS analysis (see *Methods* for more detail). (E) A representative western blot image of SA-HRP blot of streptavidin pulldown proteins from biotin untreated (-) and treated (+) Col-0, *pTNI::TbID-Myc,* and *pTNI::TbID-TNI-Myc* seedlings respectively. Experiment was repeated two times and similar results obtained. Blue and red asterisk corresponds to TNI-*TbID* and *TbID* bands respectively. Numbers on the right indicate molecular-weight markers. F) Volcano plot shows enriched proteins in biotin treated *TbID*-TNI (TbID-TNI T) Vs biotin treated Col-0 (Col-0 T) pulldown. Blue circles; enriched proteins, red circle; *TbID*-TNI, (G)TNI interactors identified through IP-MS and AP-MS are classified according to their subcellular localization. PM-Plasma Membrane; ER-Endoplasmic Reticulum.

**Table 1.**
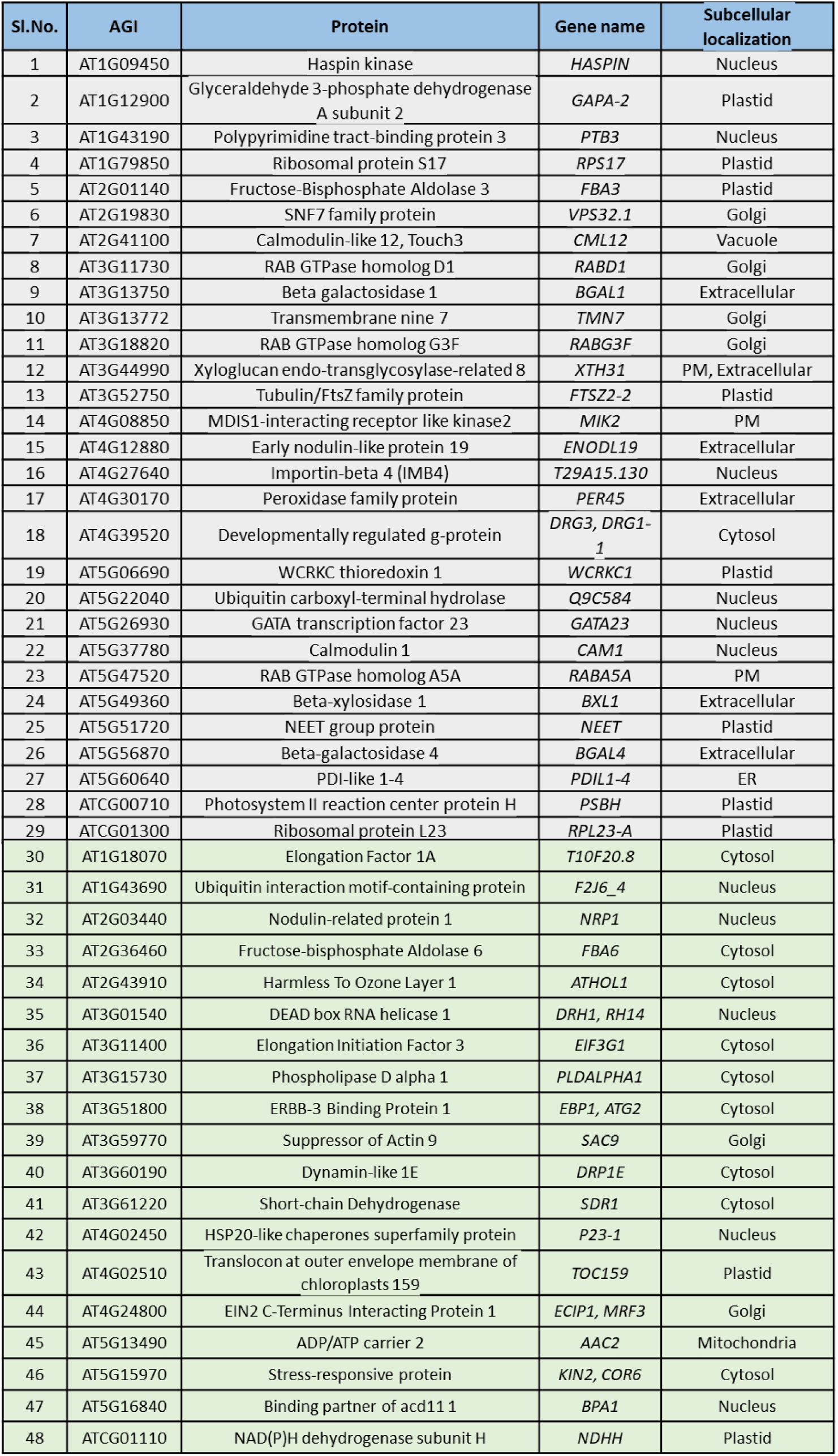
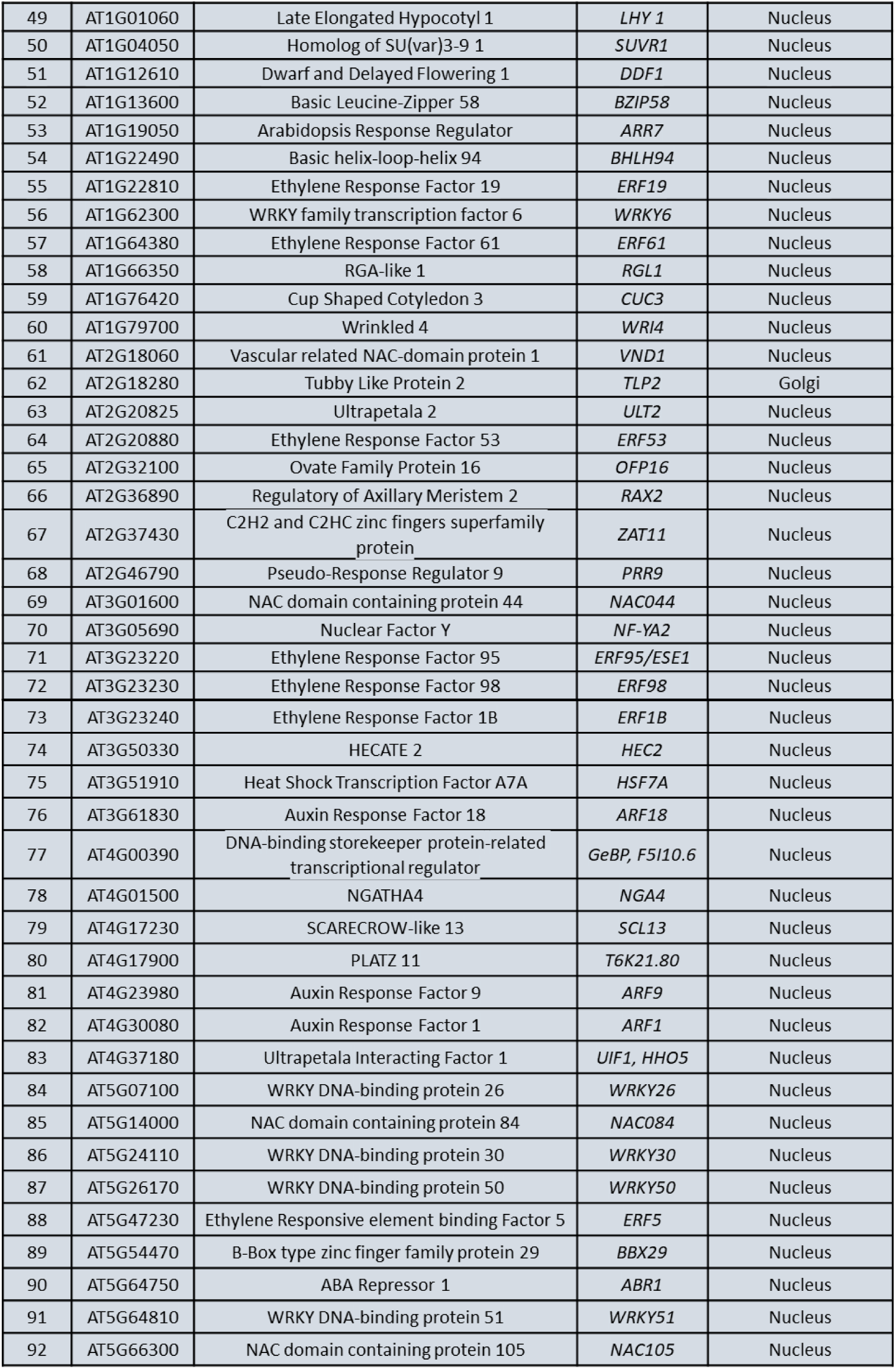
A comprehensive list of TNI-interacting proteins. TNI-interacting proteins identified in IP-MS (1-29), AP-MS (30-50), and yeast two-hybrid screen (51-95) are listed. Information on subcellular localization are from *SUBA5* database. AGI, Arabidopsis gene identifier in *TAIR* database; PM, plasma membrane, ER, endoplasmic reticulum.

The proteins identified at least two times in a group were chosen, and an enrichment graph was plotted for sample and control with changes in various parameters. In the final list of 29 proteins, 28 were selected, showing enrichment with FDR=0.05, S=2.0, and one with FDR=0.01, S=0.1. TNI profile plot showed an increase in LFQ intensity values in the samples compared to the controls. The identified 29 proteins showed a similar pattern to TNI in their respective LFQ intensity profile plot (Supp Fig. 2A).

#### **b.** Proximity labelling affinity Purification-MS (AP-MS)

We generated *pTNI::TbID-Myc and pTNI::TbID-TNI-Myc* Arabidopsis transgenic lines to perform proximity labelling and pulldown experiments. The wild type, *pTNI::TbID-Myc,* and *pTNI::TbID-TNI-Myc* seedlings were treated with the biotin. TbID is a biotin ligase enzyme that covalently adds biotin to the proteins within the radius of 10 nm (Roux et al., 2012). Seedlings without biotin treatment were taken as a negative control (Fig. 2D). We confirmed the expression of TbID-TNI (∼150kDa) and TbID (∼60kDa) with 50 µM biotin treatment (Supp Fig. 1C&D) using anti-Myc antibody. After confirming the transgenic protein expression, we standardised the biotin treatment. Based on the previous literature (Mair et al., 2019), we selected concentrations ranging from 0-200 µM. We saw a slight increase in the total biotinylated protein from 20 µM to 50 µM biotin concentration and from 30 minutes to 1 hour; it remained stable afterwards. For better enrichment in the mass spectrometry, we treated seedlings for 3 hours. We treated 5, 6, 8, and 10-day-old seedlings to check for any stage-specific increase in biotinylation and found no difference in biotinylation with the age of seedlings (Supp Fig 3A to F). Hence, we selected eight-day-old seedlings and treatment for 3 hours with 50 µM biotin concentration for further experiments. We confirmed the pulldown of biotinylated proteins by SA-HRP blotting (Fig 2E).

**Figure 3.**
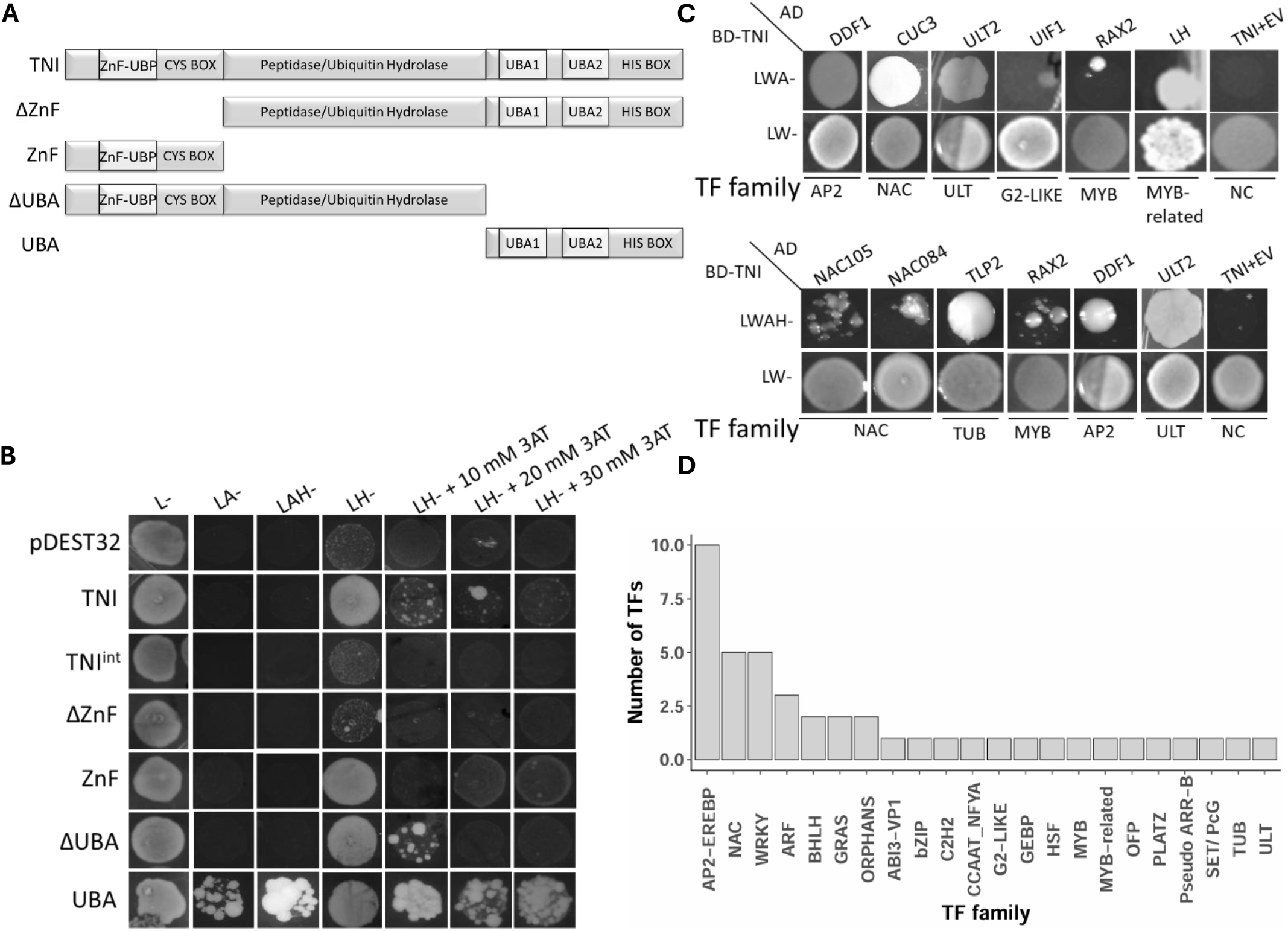
Yeast two-hybrid screen to identify TNI-interacting transcription factor (TF) proteins. (A) Schematics of full-length TNI protein and its truncated forms (ΔZnF, ZnF, ΔUBA, UBA) used as baits in yeast two-hybrid screen. ZnF-UBP, Zinc finger-ubiquitin protease domain; UBA, ubiquitin-associated domain; CYS-BOX, cysteine box domain; HIS-BOX, histidine-box domain. (B) Autoactivation assay is performed for full-length TNI and its truncated forms mentioned in A. L, A & H indicate leucine, adenine & histidine, respectively; - indicates negative, AT indicates 3-amino-1,2,4-triazole. (C) Representative images of yeast growth assay on selection plates after co-transformation with bait and prey constructs. Growth on LWA-plates signify interaction. Complete list of TFs showing interaction is shown in Table 1. BD, binding domain; AD, activation domain. (D) TFs identified in (C) as TNI-interacting proteins are categorized in different functional families.

We found 19 interacting proteins from the final list of treated to untreated controls (Fig 2F and Table 1). The enriched proteins in each combination between six samples above FDR=0.05 and S=2.0 were selected, and the proteins enriched only in the sample set were considered interactors. LFQ intensity and heat map of identified proteins across the samples and controls are plotted (Supp Fig 4&5). Further, we categorized TNI interactors from IP-MS and AP-MS according to their subcellular localization. We found that most interactors are nuclear-localized proteins, followed by cytoplasm, plastid, Golgi, and extracellular proteins, respectively (Fig 2G).

**Figure 4.**
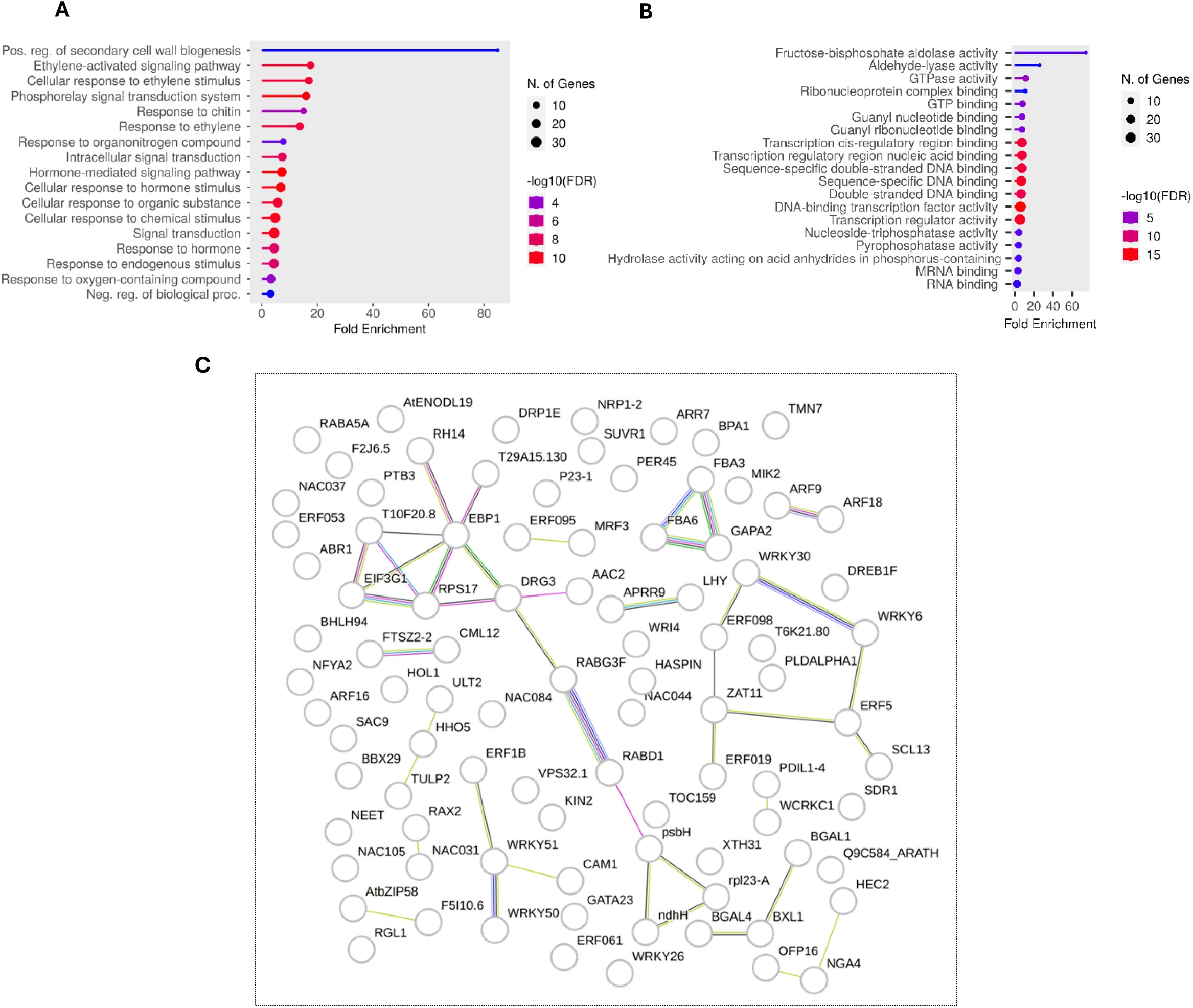
Gene ontology (GO) analysis of the TNI-interacting proteins. GO analysis performed according to biological processes (A) and molecular function (B). (C) An interaction network of all TNI-interacting proteins obtained from the STRING database. Colour of the edge depicts the method of protein identification (known interactions, pink; experimentally determined interactions from curated databases, navy blue; predicted interactions, green; gene neighbourhood, red; gene fusion, blue; gene co-occurrence, light green; text mining, black; co-expression, light blue; protein homology.

#### **c.** Yeast Two-Hybrid Assay

We saw TNI-GFP localization in the cytoplasm and nucleus (Fig. 1C). Also, previously, it was reported that TNI localizes to the nucleus (Xu et al., 2016). Since the IP-MS and AP-MS targeted overall proteins, we were interested in specifically studying TNI interaction with the nuclear-localized transcription factors. The bait vector was selected on *Leu^–^* selection in the AH109 strain with *His^–^Ade^–^* selection. Initially, we checked the auto-activation assay with full-length TNI. We observed autoactivation with *pDEST32*-*TNI* in *Leu^–^His*^–^ and *Leu^–^His*^–^ + 10-30 mM 3-aminotriazole (3-AT) conditions but not in the *Leu^–^Ade^–^* and *Leu^–^His^–^Ade^–^* conditions (Fig. 3 A). Hence, we decided to check auto-activation with truncated versions of TNI, namely *pDEST32-ΔZnF, pDEST32-ZnF, pDEST32-ΔUBA*, and *pDEST32-UBA* (Fig. 3 A). ZnF, ΔZnF, and ΔUBA showed autoactivation in the *Leu^–^His*^–^ and *Leu^–^His*^–^ + 3-AT conditions but not in the *Leu^–^Ade^–^* and *Leu^–^His^–^Ade^–^* conditions (Fig. 3 B). UBA domain showed autoactivation in all tested conditions (Fig. 3 B). Hence, we chose to perform the yeast-two hybrid screening with full-length TNI and screened for positive interactors on *Leu^–^Ade^–^* and *Leu^–^His^–^Ade^–^* plates as autoactivation was not observed under these selection conditions. (Hellmann & Estelle, 2002).We selected 197 transcription factors from the Arabidopsis prey protein library with *Trp^–^* selection to perform a yeast two-hybrid screen using TNI as bait based on the following criteria: (i) transcription factors regulating plant signalling pathways which are degraded by the 26S proteasomal system (Ramos et al., 2022; Carlos et al., 2000; Hellmann & Estelle, 2002;Li et al., 2017); (ii) Arabidopsis homologs of the *M. oryazae* transcription factors that interacted with the UBP14 homologue of the *M. oryazae* (Wang et al., 2018); and (iii) predicted TNI-interactors from the STRING database (https://string-db.org). Among the 197 transcription factors tested, a total of 44 interacted with TNI on the *Trp^–^Leu^–^Ade^–^* and *Trp^–^Leu^–^His^–^Ade^–^* selection plates (Table 1). A few representatives from these interactors are shown in Fig. 3C. We grouped the interactors according to the transcription factor family they belong to. The maximum interactors were from the plant-specific AP2-EREBP, WRKY, and NAC transcription factor families (Fig. 3D).

### GO analysis

To understand the biological processes involved by the interacting partners of TNI, we performed a gene ontology (GO) analysis of the interactors obtained from all 3 approaches. Major biological processes associated with the interactors were hormone signalling, response to environmental stimuli, signal transduction, secondary cell wall biogenesis and molecular functions like carbohydrate metabolism, nucleotide binding, and hydrolase activity (Fig. 4 A, B). Some of the protein mutants showed phenotypic similarity with the *tni* mutant, and they are reported to function in a complex as predicted or experimentally confirmed in the STRING database (Fig. 4C). The flower petal regulation proteins ULTRAPETALA2 (ULT2), ULTRAPETALA1 INTERACTING FACTOR1 (UIF1), and TUBBY-LIKE PROTEIN 2 (TLP2) are found by text-mining. Co-expression of circadian clock proteins LATE ELONGATED HYPOCOTYL (LHY) and PSEUDO RESPONSE REGULATOR (PRR9); cell wall remodellers BETA-GALACTOSIDASE 1, BETA-XYLOSIDASE 1, BETA-GALACTOSIDASE 4 are reported from the curated databases (Arias et al., 2014; Lee et al., 2007). The interaction of DEVELOPMENTALLY-REGULATED G-PROTEIN 3 (DBG3) and ADP, ATP carrier protein 2 (AAC2); AUXIN RESPONSE FACTOR 9 (ARF9) and AUXIN RESPONSE FACTOR 18 (ARF18) have been experimentally confirmed in *Saccharomyces cerevisiae* (Costanzo et al., 2016; Trigg et al., 2017)(Fig 4C). This suggests that TNI might be directly or indirectly interacting with the protein complex to regulate cellular activities.

### ULT2 interacts with TNI

The *ult1-2 ult2-3* double mutant shows an increase in petal number similar to the *tni* mutant. ULT2 regulates flower petal numbers along with another member of the family, ULT1 (Monfared et al., 2013). Probably, TNI interaction stabilises ULT2 to regulate the petal number. We validated the interaction of the ULT2 from the Y2H screen and TNI by bimolecular fluorescence complementation (BiFC) assay. We observed that ULT2-nEYFP interacts with the TNI-cEYFP upon co-expression in the Nicotiana leaf (Fig 6). We found an interaction of TNI with UIF1, another TF involved in regulating flower petal numbers in the Y2H screen. UIF1 interacts with ULT1, and ULT1 interacts with ULT2 (Monfared et al., 2013; Moreau et al., 2016). It is also possible that ULT1, ULT2, and UIF1 might be interacting in a complex with TNI.

### HASPIN kinase is stabilized in the *tni* mutant

We selected an interactor identified in our experiments to further investigate the physiological and biochemical implications of its interaction with TNI. We chose haspin kinase, an interactor obtained in the IP-MS approach with the highest LFQ intensity, suggesting its strong interaction with the TNI (Supp Fig 2A). Haspin kinase was shown to phosphorylate Histone 3 at Thr3 and Thr11 both *in vitro* and *in vivo* (Karimi-Ashtiyani et al., 2013; Kurihara et al., 2011). The downregulation of haspin resulted in multiple shoots, abnormal flowers, semi-fertile and undeveloped siliques, whereas the overexpression lines showed multi-rosette and adventitious shoot apical meristems (Karimi-Ashtiyani et al., 2013). Since the *tni* mutant shows several related phenotypes, such as increased petal number, decreased shoots, undeveloped siliques, and reduced seed number(Karidas et al., 2015; Majumdar and Karidas et al., 2020), we examined the levels of haspin protein in the *tni* mutant. For this purpose, we generated a *pHaspin::Haspin-sGFP* lines and crossed it with *tni* mutant to obtain *pHaspin::Haspin- sGFP*tni* lines. Using confocal microscopy, we could detect an increased Haspin-GFP signal in the *pHaspin::Haspin-sGFP #1*tni* line as compared to the *pHaspin::Haspin-sGFP #1* (Fig 5A&B). To confirm this further, we performed western blotting using total protein extracts from two independent lines of *pHaspin::Haspin-sGFP* and their respective crosses with *tni* mutant. Haspin-sGFP band was detected in *pHaspin::Haspin-sGFP #1*tni* but not in the *pHaspin::Haspin-sGFP #1*. Together, these observations indicated that TNI negatively regulates the protein abundance of Haspin kinase. (Fig 5C). To investigate the genetic interaction of *HASPIN* with *TNI*, we crossed several RNAi-based knockdown lines and constitutive overexpression lines with the *tni* mutant (Karimi-Ashtiyani et al., 2013).We found a significant reduction in the number of seeds per silique in *HaspinRNAi #23.2* and *HaspinRNAi #14.1* compared to Col-0. While *HaspinRNAi #23.2*tni* and *HaspinRNAi #25.1*tni* showed a seed number similar to *tni*, the *HaspinRNAi #14.1*tni* exhibited a significantly reduced seed number than *tni* (Fig 5D). The overexpression lines of haspin, namely *35S::Haspin #2.1* and *35S::Haspin #10.2,* showed a decreased seed number than Col- 0, whereas the seed number in *35S::Haspin #2.1*tni* and *35S::Haspin #10.2*tni* was similar to *tni* mutant (Fig 5E). The reason might be that wild-type TNI in the *tni* mutant is probably destabilizing haspin or the involvement of other pathways. A further decrease in the seed number in *HaspinRNAi #14.1*tni* as compared to both parental lines indicate the possibility of other parallel pathways through which these factors regulate seed number.

**Figure 5.**
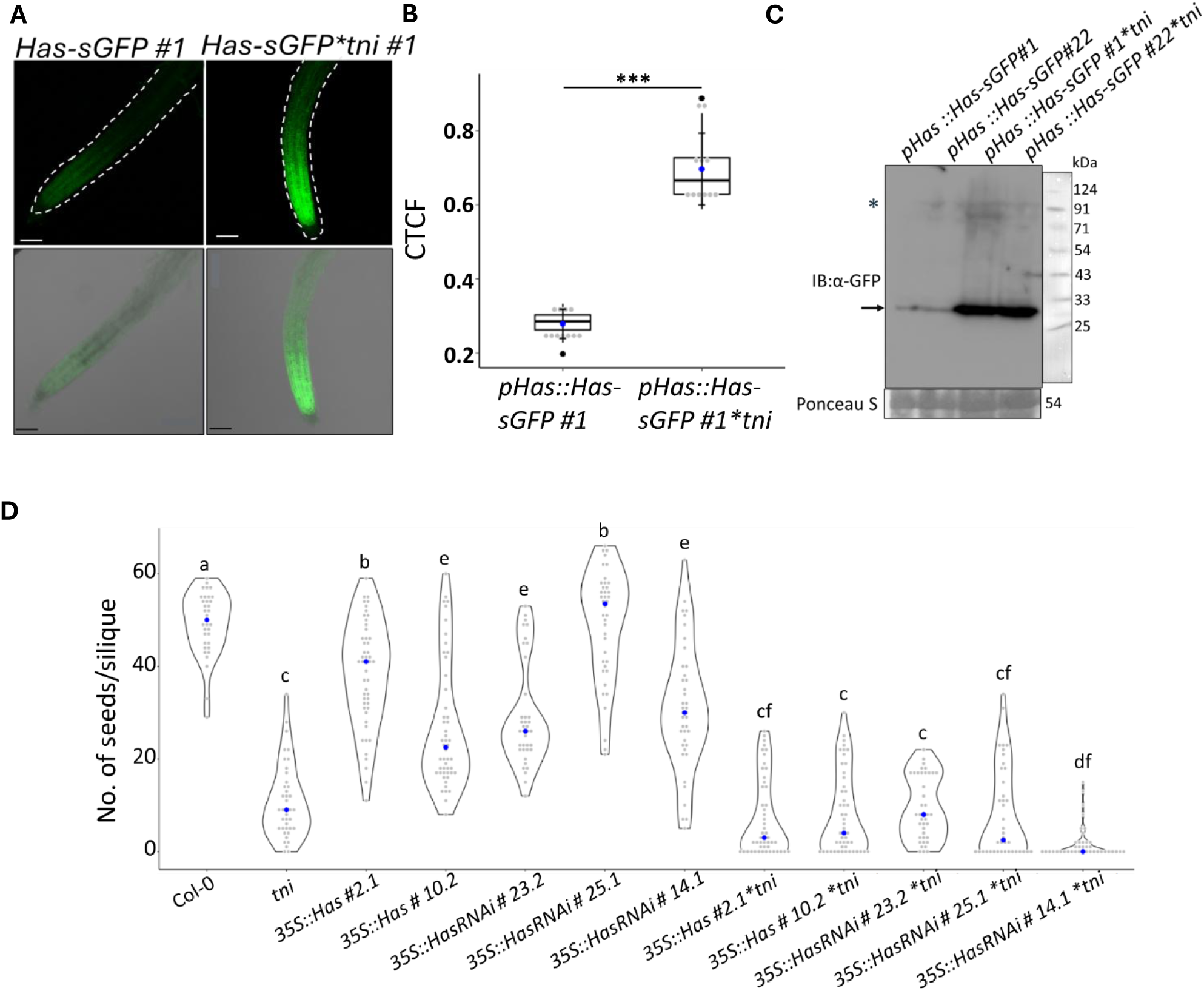
Haspin kinase is stabilised in the *tni* mutant. (A, B) Images of root tips (A) of 8-day old *pHaspin::Haspin-GFP* and *pHaspin::Haspin-GFP*tni* seedlings showing GFP fluorescence (#1 indicates line number), and average quantification (n=11) of fluorescent intensity (B). CTCF indicates Corrected Cell Fluorescence Intensity. Error bars represent the SD; asterisk indicates statistical significance using unpaired Student’s *t* test (*** *p* = 0). Scale bar in (A), 100 µm. (C) Image of western blot of total protein extracted from 8-day old seedlings of indicated genotypes and probed with α-GFP antibody. Asterisk and arrow correspond to Haspin-sGFP protein band and cleaved GFP protein, respectively. Ponceau staining shown below the western blot served as the loading control. Numbers on the right indicate molecular weight. (D) Average number of seeds per silique (n = 35-40) in the indicated genotypes. Numbers on the right indicate independent transgenic events. Each grey dot indicates number of seeds in a silique. Blue circle represents the median. Groups indicated by the same letter are not statistically different according to ANOVA with Tukey’s HSD test.

**Figure 6.**
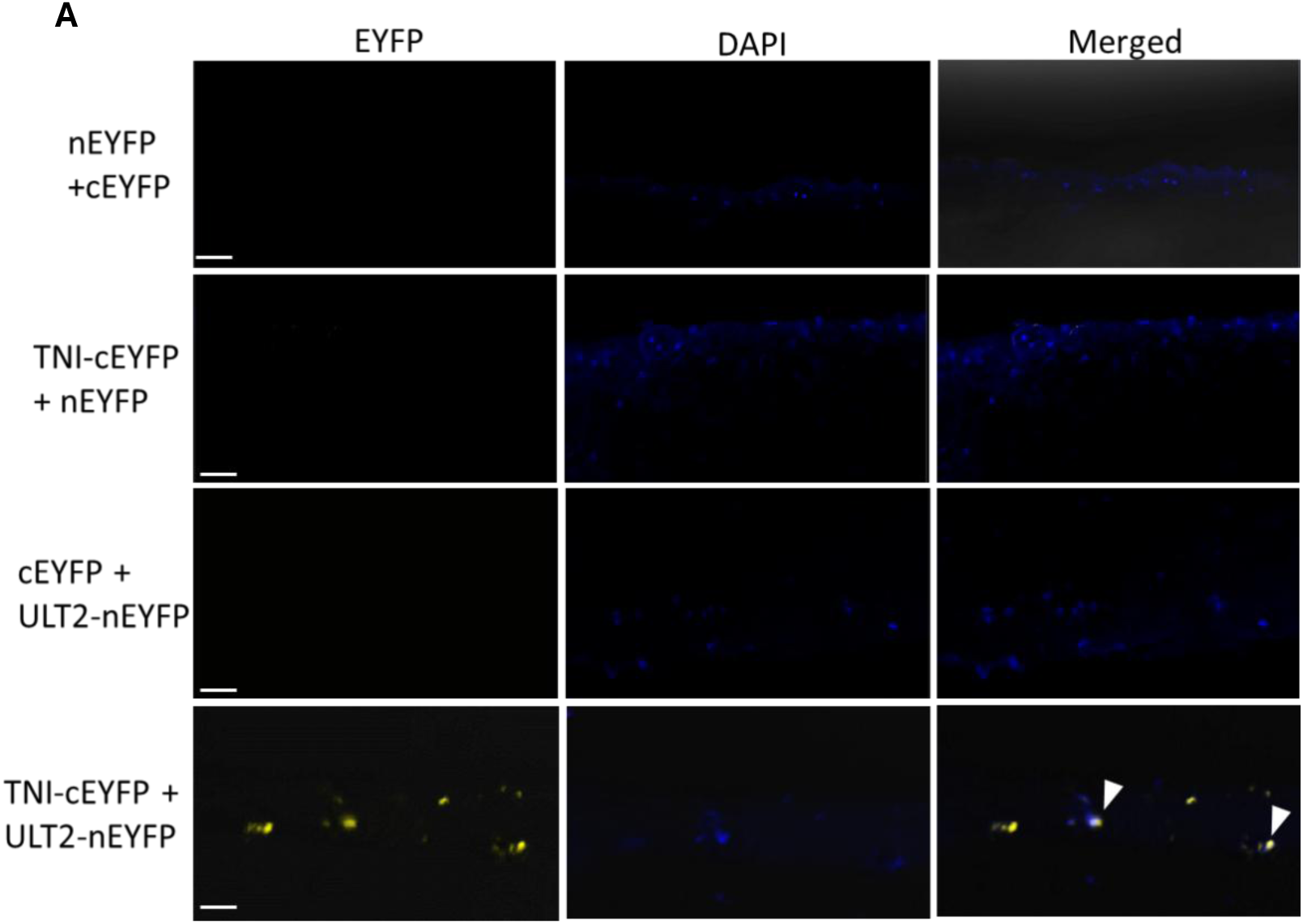
*In vivo* interaction of TNI with ULT2. Bimolecular fluorescence complementation assay of TNI-cEYFP and ULT2-nEYFP transiently expressed in tobacco leaves. Images of the abaxial epidermal cells were taken 48-72 hrs post-infiltration. EYFP fluorescence, DAPI fluorescence, and merged images are shown. Images in the first three rows serve as negative controls. The fourth lane shows co-infiltrated test proteins TNI-cEYFP and ULT2-nEYFP proteins interacting in the nucleus (white arrowhead). Scale bar, 50 µm.

## Discussion

We used three different approaches to identify a broader TNI interactome that would generate a fair understanding of the role of TNI/UBP14 in plant development. TNI/UBP14 is present throughout all stages of growth and in all tissues. TNI/UBP14 is essential in maintaining plant protein homeostasis. The absence of TNI/UBP14 causes embryonic lethality, and its hypomorphic alleles are recessive (Karidas et al., 2015; Tzafrir et al., 2002; Xu et al., 2016). This suggests the importance of the TNI/UBP14 in the UPS system. It has been shown that TNI/UBP14 can bind to the free polyubiquitin chain and disassemble it (Majumdar and Karidas et al., 2020). It is likely that TNI/ UBP14 also cleaves the ubiquitin moiety from target proteins as well. Our study aimed to identify those target proteins by identifying a TNI-interacting subset of the proteome.

We found proteins involved in diverse biological processes from the three approaches used here, including cell wall remodelling, stress response, carbohydrate metabolism, protein synthesis, chaperones, histone-associated proteins and epigenetic modifiers. Interestingly, nearly half of the interacting proteins are predicted to function as enzymes, suggesting the role of TNI/UBP14 in diverse biological processes in the cell. Several proteins exhibit functional similarity among the interactors and are grouped under the same pathway.

An intriguing outcome of our study is the lack of overlapping interactors among the subsets identified in various methods. The non-overlapping targets perhaps indicate that a more inclusive approach should be taken in future to identify the full repertoire of the interacting proteins of a specific bait. It has been demonstrated that the same method used for identifying interacting proteins sometimes identifies targets with minimal overlap. For example, a comparative bioinformatic analysis found merely ∼20% overlapping proteins from the union of three large-scale Y2H screenings aimed to identify protein-protein interaction in the yeast (Yu et al., 2008).

Since we were interested in building a comprehensive interactome of TNI/UBP14, employing approaches with different principles best suited our purpose. The yeast two-hybrid study, the IP-MS study, and the AP-MS protocol fundamentally differ in their approach and, hence, explain the outcome of non-overlapping target proteins from these experiments. Y2H study is an out-of-context, condition-specific investigation focussing on interaction upon nuclear localization and tends to show results with transient interactors. *In vivo* approaches such as IP-MS detect stable interactors, whereas AP-MS identifies proteins that make stable complexes with the bait and those that come in proximity to the bait without even making any direct contact (Roux et al., 2012).

As TNI/UBP14 is a ubiquitin protease, we examined whether any identified interactors overlap with the previously reported ubiquitinated proteins. We compared the TNI/UBP14 interactors with the ubiquitinated proteins database identified through the pulldown approach. We identified that thirty-two proteins from our dataset are ubiquitinated, as reported in the two previous studies (Kim et al., 2013; Song et al., 2021)(Fig. Supp 6 A, B). This suggests that the TNI-interacting proteins perhaps are degraded by 26S proteasomal machinery. However, the ubiquitination status of the target proteins identified in our study should be compared between wildtype and *tni* mutant plants, and this remains a major limitation of our work.

When we analysed all our interacting proteins in the STRING database, we saw that protein complexes performing specific functions were predicted or confirmed earlier. The largest complex we identified comprised eleven protein nodes with ten edges involved in protein synthesis. We also found that several target proteins are co-expressed with TNI, including BETA-GALACTOSIDASE 1, BETA-XYLOSIDASE 1, BETA-GALACTOSIDASE 4 involved in cell wall remodelling, ULT2, UIF1, and TLP 2 involved in regulating flower petal number, REGULATOR OF AXILLARY MERISTEM 2 (RAX2), and NO APICAL MERISTEM 031 (NAC031) involved in axillary branching. Physical interaction of FILAMENT TEMPERATURE-SENSITIVE MUTANT 2 (FTSZ2), CALMODULIN 12 (CML12) and, FRUCTOSE BISPHOSPHATE ALDOLASE 3 (FBA3), FBA6, and GLYCERALDEHYDE 3 PHOSPHATE DEHYDROGENASE A SUBUNIT 2 (GAPA2), which appeared in STRING analysis, is well reported (Fig. 3C).

We have validated the interaction of the selected interactors by other binary methods. Haspin stabilization in the *tni* mutant suggests that TNI/UBP14 is required for the degradation of haspin in the wild type. Considering the broad range interactome of TNI/UBP14 and the pleiotropic phenotype of its mutant (Karidas et al., 2015; Xu et al., 2016)it would be interesting to examine the biological significance of TNI interaction with these diverse interactors. An in-depth study on the affinity of TNI towards ubiquitinated or non-ubiquitinated states is necessary to understand the biological function of these interactions in plants.

## Materials and Methods

### Plant materials and growth conditions

Seeds were surface sterilized and stratified in darkness for three days at 4°C; they were transferred to the growth chamber and kept in light for 6-9 hours. Later, plates were kept under dark conditions for three days to select the transformants in hygromycin selection and shifted back to light. Plants were maintained under long-day conditions with 16 h light (120 μmol/m^2^/s) and 8 h dark, 22°C. All transgenic lines were generated in the Col-0 background. To perform the TNI protein pulldown experiments, seedlings were sown on half MS medium and stratified in the dark at 4°C for two to three days. Later, plates were shifted to light and maintained under the long-day condition with 16 h light (120 μmol/m^2^/s) and 8 h dark, 22°C.

### Construct generation and plant transformation

The TNI CDS (2394 bp) was subcloned by amplifying from the *TNI-pGEX4T1* using CDS- specific primers with 5’ attB1 and 3’attB2 sites and cloned into the entry vector pDONOR221 by BP Clonase reaction (Invitrogen) (Prof. Imran Siddique acknowledged). TNI promoter was amplified from the *pTNI-pCAMBIA* (Premananda Karidas, PhD thesis, 2014) using promoter-specific primers with 5’ attB4 and 3’attB1R sites, respectively, and cloned by pENTR^TM^ 5’- TOPO^TM^ TA Cloning^TM^ kit (Invitrogen). Turbo ID was amplified from the vector *pVHS64* (Dr. P.V. Shivaprasad acknowledged), and it is cloned to the N-terminus end of the TNI in the vector *TNI-pGEX4T1* by Gibson Assembly Master Mix (E2611S, NEB). *TbID -TNI* was amplified by CDS-specific primers with 5’attB1 & 3’attB2 sites and cloned into the pDNR221 vector by the BP clonase reaction (Invitrogen). 35S promoter was amplified from the pCAMBIA1301, and the RPS5A promoter was amplified from the plasmid pRPS5A_ER8_GW (Dr. Ram Yadav & Dr. Nam Hai Chua acknowledged) using promoter-specific primers with 5’ attB4 and 3’attB1R site respectively and cloned into the entry vector pDONOR P4-P1R by BP Clonase reaction (Invitrogen). Further, promoter and CDS entry clones are incorporated into the R4pGWB vectors (spectinomycin selection) with the C-terminal reporter tags R4pGWB 510-FLAG, R4pGWB 519-10xMyc, R4pGWB 533-GUS, R4pGWB 543-G3GFP by tripartite LR reaction (Invitrogen), transformed into the *Agrobacterium tumifaciens* GV3101 strain by electroporation (Biorad, 2.5Hz, 5 seconds). Positive clones were selected, and floral dipping was performed in Col-0 plants. Transformants were selected on MS medium (Himedia) containing 100 µg ml^-1^ cefotaxime sodium salt (Himedia) and 20 µg ml^-1^ hygromycin (Sigma-Aldrich). *pHaspin::Haspin-sGFP* construct was obtained and the above procedure was followed to generate transgenics (Prof. Daisuke Kurihara acknowledged). The selected transgenic lines were crossed to the *tni* mutant, and positive plants were used for the biochemical experiment. All *35S::Haspin* and *HaspinRNAi* lines were crossed to *tni* mutant (Prof. Andreas Houben acknowledged).

### Construct generation and Yeast two-hybrid assay

TNI CDS from *pDONOR221-TNI* (see above) was moved C-terminal to the GAL4 DNA-binding domain (GAL4-DBD) in pDEST32 vector by LR reaction (Invitrogen) and introduced into yeast strain AH109 by the LiOAc/ssDNA/PEG method (Gietz & Schiestl, 2008). Transformants were selected on a synthetic drop-out medium lacking leucine. A prey library of 1956 Arabidopsis transcription factors C-terminal to the GAL4 DNA-activation domain (GAL4-AD) in the vector pDEST22 was received in the form of bacterial glycerol stocks from the ABRC stock centre (Pruneda-Paz et al., 2014). Plasmid isolation was carried out from the complete library, and in those, 197 plasmids were transformed individually into the previously transformed AH109 yeast strain expressing TNI by the standard LiAc protocol and grown on *Leu^–^His^–^* +3AT, *Leu^–^Ade^–^, Leu^–^His^–^ Ade^–^* selection as described in the Yeast Protocols Handbook (Clontech, http://www.clontech.com)

### Immunoblot -analysis

Protein extraction was carried out from 29-30-day-old leaves using protein extraction buffer (50 mM Tris-HCl pH 7.4, 150 mM NaCl, 10 mM MgCl2, 0.1% (v/v) NP-40,1 mM PMSF, and 1X complete protease inhibitor cocktail (Roche)). The crude extract was centrifuged at 12,000 g for 15 min at 4°C. Protein concentration was measured by Bradford assay, and equal concentrations of protein were loaded into the 10% (v/v) SDS-PAGE, transferred to PVDF membrane (Millipore, USA), and immunoblotted using anti-MYC (#2276, CST), and anti-GFP (#3H9, Chromotek) primary antibodies and anti-Mouse (Sigma), and anti-rat secondary antibodies. ECl (Millipore, USA) was used to develop the blots. For the biotin-treated seedlings, the PVDF membrane was blocked in 5% BSA and incubated in streptavidin-HRP (S911, Thermo Fisher Scientific) for 1 hour, and proceeded to develop.

### Bimolecular Fluorescence Complementation assay

*pDONOR221-TNI* and *pDONOR221-ULT2* was cloned into the *pSITE-cEYFP-N1* and *pSITE nEYFP-*C1 by LR reaction respectively. Further, these clones were transformed into the Agrobacterium tumifaciens GV3101 strain by electroporation, including the empty BiFC binary vectors. (Biorad, USA, 2.5Hz, 5 seconds). Positive clones were selected, and the primary culture was grown overnight with respective antibiotic selection. Inoculum from the primary culture was used for the secondary culture for four hours of growth at 30°C. Cells were pelleted down at 4000rpm, 20 minutes at 4°C. The pellet was resuspended in the MES buffer (100 mM MES-KOH, pH 5.6, 10 mM MgCl2, 15 mM Acetosyringone) to OD600=0.8. The respective cultures were mixed in equal volumes and incubated on ice for one hour. The cultures were infiltrated onto the abaxial side of the 3–4-week-old Nicotiana leaves and plants were watered well and kept in the dark for 24 hours. Then, the plants were shifted to light for 24-36 hours, and the leaves were DAPI stained and observed under the confocal microscope with EYFP (514 nm) and DAPI (358 nm) 405 diode, Argon multiline laser under 10X magnification (Zeiss 880 Multiphoton, Germany).

### Immunoprecipitation-pulldown

The eight-day-old *RPS5a::TNI-Myc* seedlings grown in half MS medium were treated in 20 µM MG132 for 16hrs at 22°C. Seedlings were tap-dried, snap-frozen, and stored at -80°C. Seedlings were ground in IP buffer I (50 mM HEPES pH 7.4, 50 mM NaCl,10 mM EDTA, 0.1% TritonX-100, 1 mM PMSF, 0.1 mg/ml Dextran, 1X complete protease inhibitor cocktail (Roche), 50 µM MG132) and centrifuged at 12,000rpm for 20 minutes at 4°C. The supernatant was separated, and protein concentration was measured by the Bradford method. Protein extract from *RPS5a::TNI-Myc* seedlings were incubated with anti-Myc magnetic beads (#5698S, CST) and anti-IgG mouse (#5873S, CST) at 4°C on a rotor for 5hrs. Beads were separated by a magnetic stand and washed twice in IP buffer II (50 mM HEPES pH 7.4, 50 mM NaCl, 10 mM EDTA), once in IP buffer III (50 mM HEPES pH 7.4, 150 mM NaCl,10 mM EDTA) and once in 1X PBS, pH 7.4. The buffer was removed entirely from the beads and stored at -80^°^C until proceeded for the on-bead digestion.

### Affinity Purification by streptavidin beads

Eight-day-old Col-0, *pTNI::TbID-Myc,* and *pTNI::TbID-TNI-Myc* seedlings grown on half MS were treated with 50 µM biotin for 3hrs at 22°C, and biotinylation was stopped by washing in ice-cold water three times for three minutes each. Seedlings were tap-dried and snap-frozen, and an aliquot was taken for immunoblot analysis. Protein was extracted in extraction buffer as mentioned in Mair et al., 2019 (50 mM Tris pH 7.5, 150 mM NaCl, 1 mM EGTA, 1 mM DTT, 0.1% SDS, 1% TritonX-100, 0.5% Sodium deoxycholate, 1X complete protease inhibitor cocktail (Roche),1 mM PMSF) and kept at 4°C for 15 minutes on rotor wheel. Protein extract was centrifuged at 12,000 rpm for 15 minutes at 4°C. The supernatant was separated and depleted of extra biotin by passing through the PD-10 desalting column (GE-Healthcare). Protein concentration was measured by Bradford assay, and an equal concentration of protein was taken and incubated with Dynabeads MyOne Streptavidin C1 (#65002, Invitrogen) overnight at 4°C on the rotor. Beads were separated on a magnetic rack and washed in respective buffers, as described in Branon et al. (2018). Beads were washed with 1 ml each of the following solutions on rotor 2x with cold extraction buffer at 4°C, 1x with cold 1 M KCl at 4°C, 1x with cold 100 mM Na_2_CO_3_ at 4°C, 1X with 2 M Urea in 10 mM Tris pH 8.0 at room temperature and 2X with cold extraction buffer without complete protease inhibitor and PMSF at 4°C. 20ul or 2% of the beads were boiled in 20 ml 4x Laemmli buffer with 20 mM DTT and 2 mM biotin at 98°C for 15 min for immunoblots. Buffer was removed from the rest of the beads and stored at -80°C until further procedure.

### MS Sample Preparation

On-bead trypsin digestion was performed to digest the proteins. Beads were washed twice in 50 mM Tris pH 7.5 and 1 ml of 2 M urea in 50 mM Tris pH 7.5. Beads were incubated at 25°C for 3 hrs with shaking in 80 µl of Trypsin buffer (50 mM Tris pH 7.5, 1 M Urea, 1 mM DTT, 0.4 µg Trypsin). The supernatant was transferred to a new tube, and beads were washed twice in 60 µl of 1 M Urea in 50 mM Tris pH 7.5. Eluates were pooled and reduced by adding DTT to a final concentration of 4 mM and incubated at 25°C for 30 minutes. Alkylation was performed using 10 mM Iodoacetamide to final concentration followed by 0.5 µg Trypsin digestion overnight at 25°C with shaking. The next day, the supernatant was separated from the beads and desalted on Pierce ™ C18 Spin columns (# 89873, Thermo Scientific).

### LC-MS/MS

In the LC-MS/MS run, peptides were resuspended in 0.1% formic acid and 0.1% acetonitrile. Samples were analyzed in a PepMap^TM^ RSLC C18 Nano liquid chromatography-Orbitrap Fusion mass spectrometer (Thermo Fisher). Peptides were separated by PN ES803A rev.2 (Thermo Fisher) analytical column. The flow rate was 250 nl/min, and a 160 min gradient was used. Elution of the peptide was done in a 5% to 95% gradient of solvent B (80% acetonitrile, 0.1% formic acid, 20% water) with a corresponding gradient of solvent A (0.1% formic acid) for 160 minutes, followed by 60 min wash in solvent B. Precursor Scan Range(m/z) was 350 to 2000 and 20 topmost intense charged precursors were selected for fragmentation. Fragmentation was done by Orbitrap-higher energy collision dissociation (OT/OT/HCD) with 30% HCD collision energy.

### MS data analysis

Protein identification was performed in Max Quant (version 2.4.2.0) with minor modifications in the default settings. The parameters for the antibody-mediated pulldown experiment are as follows: Control IgG pulldown and sample anti-Myc pulldown were set as fractions 1 and 2, respectively. Under the “Group-specific parameters,” oxidation (M) and acetyl (Protein N-term) were set as variable modifications with a maximum of five modifications per peptide. ‘Digestion’ was set to Trypsin/P, with two maximum missed cleavages. The minimum ration count was set to one with Normalization Type-Classic under the ‘Label-Free Quantification.’ In the “Global parameters,” Oxidation(M) and Acetyl (Protein N-term) are set as modifications used in the protein quantification. “Unique+Razor peptides” were used for quantification. PSM FDR and Protein FDR were set to 0.01, and the Match between runs was selected under ‘Identification.’ Peptides were searched against the TAIR10 database (Araport_pep_20220103, updated on 2022-01-03, www.arabidopsis.org) with 48231 protein entries with additional MYC, trypsin and keratin as contaminants. The minimum peptide length was set to seven.

The settings were set as above, with slight changes for the enzyme-based proximity labelling approach. Fractions were set from 1-6 for the untreated and treated Col-0, Turbo, and Turbo-TNI, respectively. Carbamidomethyl(C) is under fixed modification, and Trypsin is under the ‘Digestion.’ Peptides were searched against the TAIR10 database (Araport_pep_20220103, updated on 2022-01-03, www.arabidopsis.org) with 48231 protein entries with additional MYC, and TurboID sequences as contaminants.

### Confocal imaging

The root tip of the 8 DAS of *pTNI::TNI-G3GFP*, *HASPIN-sGFP* and *HASPIN-sGFP*tni* were imaged under the confocal microscope with the GFP (395 nm), Argon multiline laser under 10X magnification (Zeiss 880 Multiphoton, Germany).

### Statistical analysis

The ‘proteinGroups.txt’ file was uploaded to Perseus (version 2.0.7.0) for the statistical analysis. T-test, Histogram, Profile plot, Heat map, and PCA analysis were performed in Perseus. Subcellular localization prediction was done in SUBA5 (https://suba.live), and results were exported to R Studio (version 4.3.0) for plotting the graph. Gene Ontology analysis was performed in Shiny GO (V0.80)(Ge et al., 2020) and STRING database (version 12.0). Graphs were plotted in R Studio (version 4.3.0).

