## supplemental file for "Identification and Characterization of Interacting Proteins of TARANI/Ubiquitin Specific Protease-14 in *Arabidopsis thaliana*"

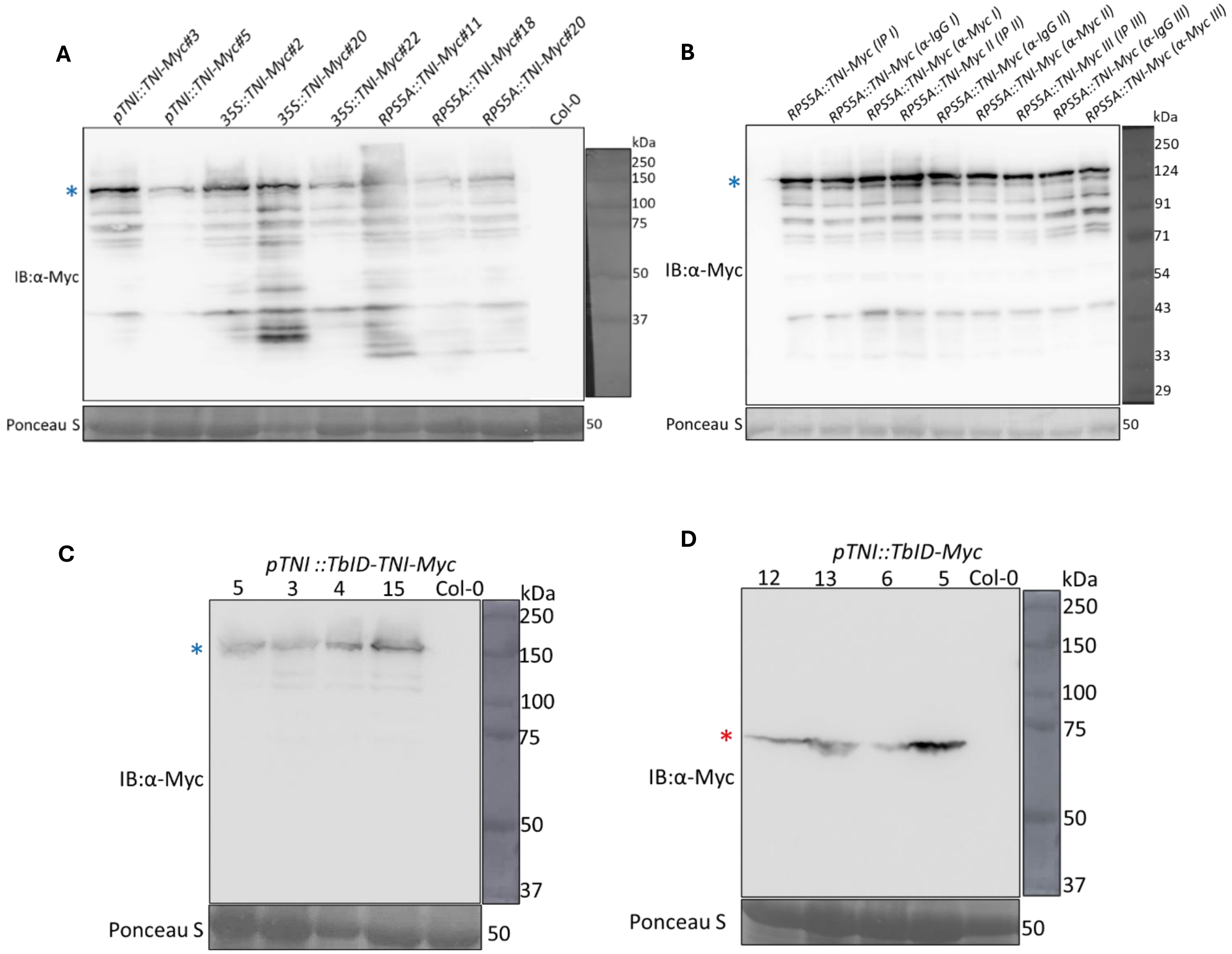


**Supp. Figure 1. Expression of recombinant proteins in the transgenic lines.** (A) Images of western blot of total protein extract from Col-0 and TNI transgenic lines using α-Myc antibody. (B) Input with total protein and flowthrough of *RPS5A::TNI-Myc* protein pulldown of three biological replicates. Blue asterisk marks the TNI-Myc protein band. (C) Col-0, *pTNI::TbID-TNI-Myc,* and (D) Col-0, *pTNI::TbID-Myc* transgenic lines (blue and red asterisk marks TbID-TNI-Myc and TbID-Myc protein bands respectively). Ponceau staining shown below the western blots served as the loading control. Numbers on the right indicate molecular weight.

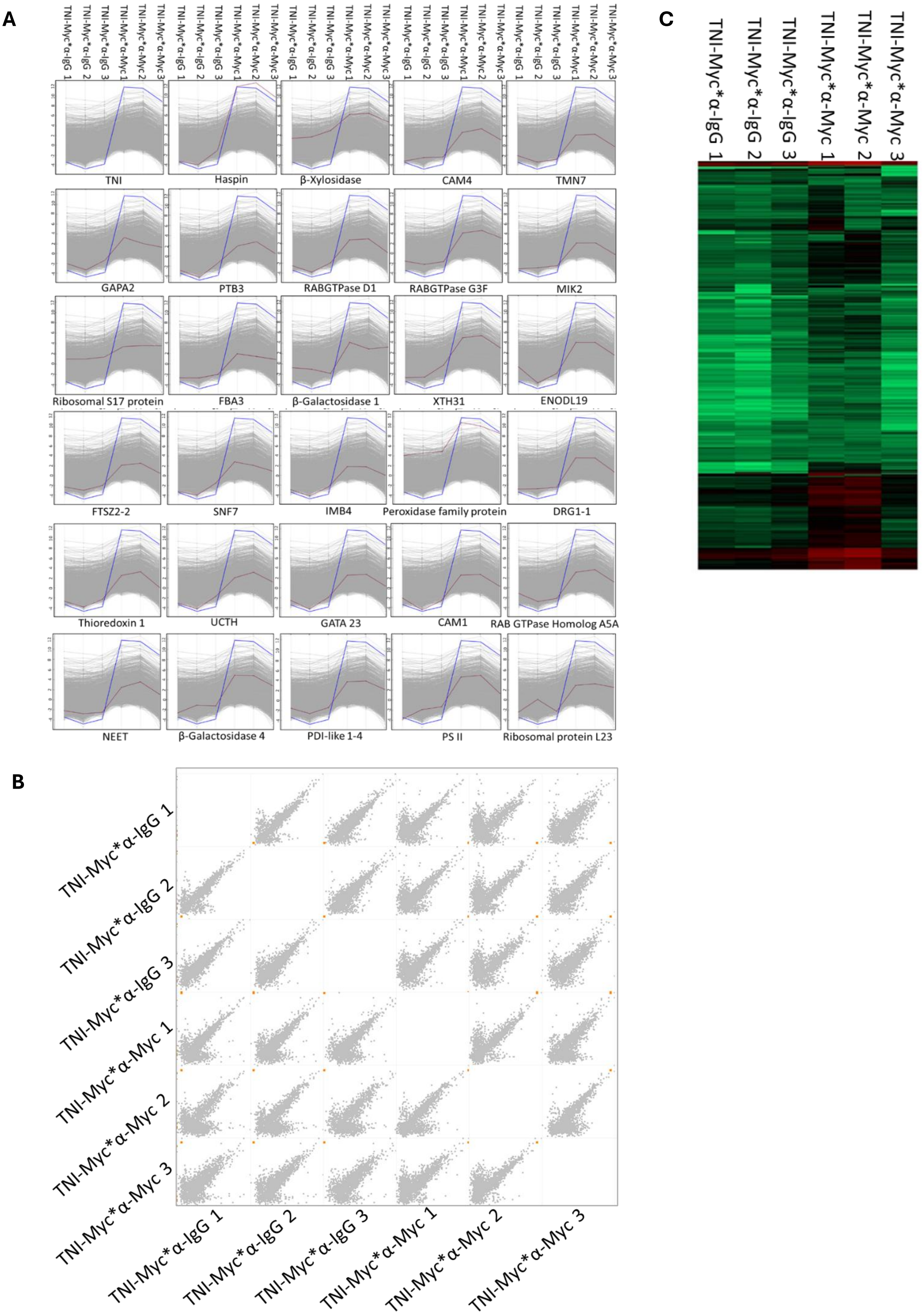


**Supp. Figure 2. Statistical analysis of TNI-interacting proteins from IP-MS**. (A) Label Free Quantification intensity profile plot and (B) multi-scatter plot, showing relationship among the samples and controls. Grey dots/ lines, interacting proteins; red dot/ line, TNI protein. (C) Heat map of all the pulldown replicates.

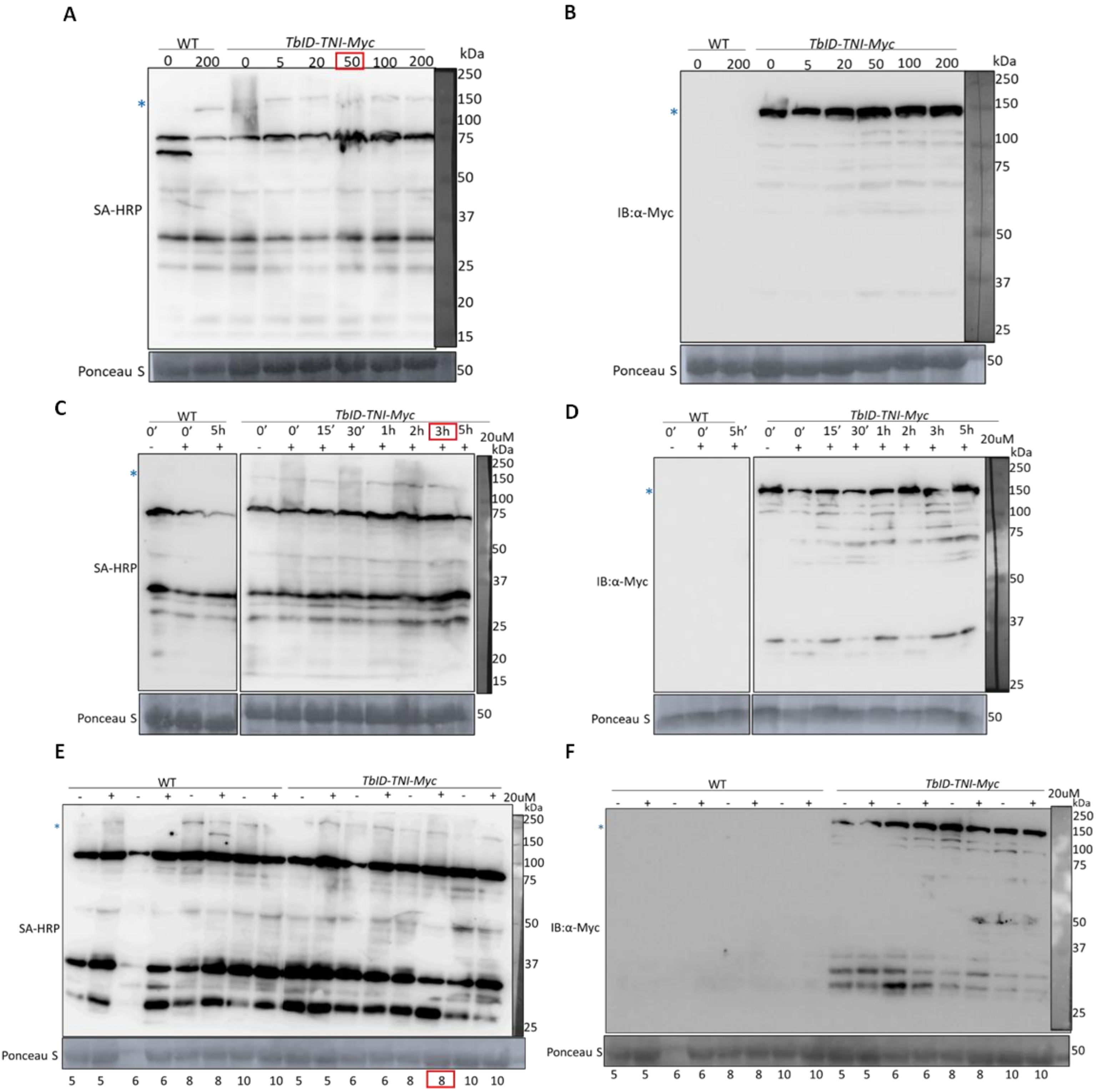


**Supp. Figure 3. Standardisation for AP-MS**. (A, C, E) Western blot images of total protein extracts from Col-0 and *pTNI::Turbo-TNI-Myc* probed with SA-HRP and (B, D, E) α-Myc antibody at different concentrations of biotin (A, B), for indicated durations (C, D), and days after stratification (E,F) Blue asterisk marks the TbID-TNI band. Red square highlights the selected condition for the pulldown-experiment. Ponceau staining shown below the western blots served as the loading control. Numbers on the right indicate molecular weight.

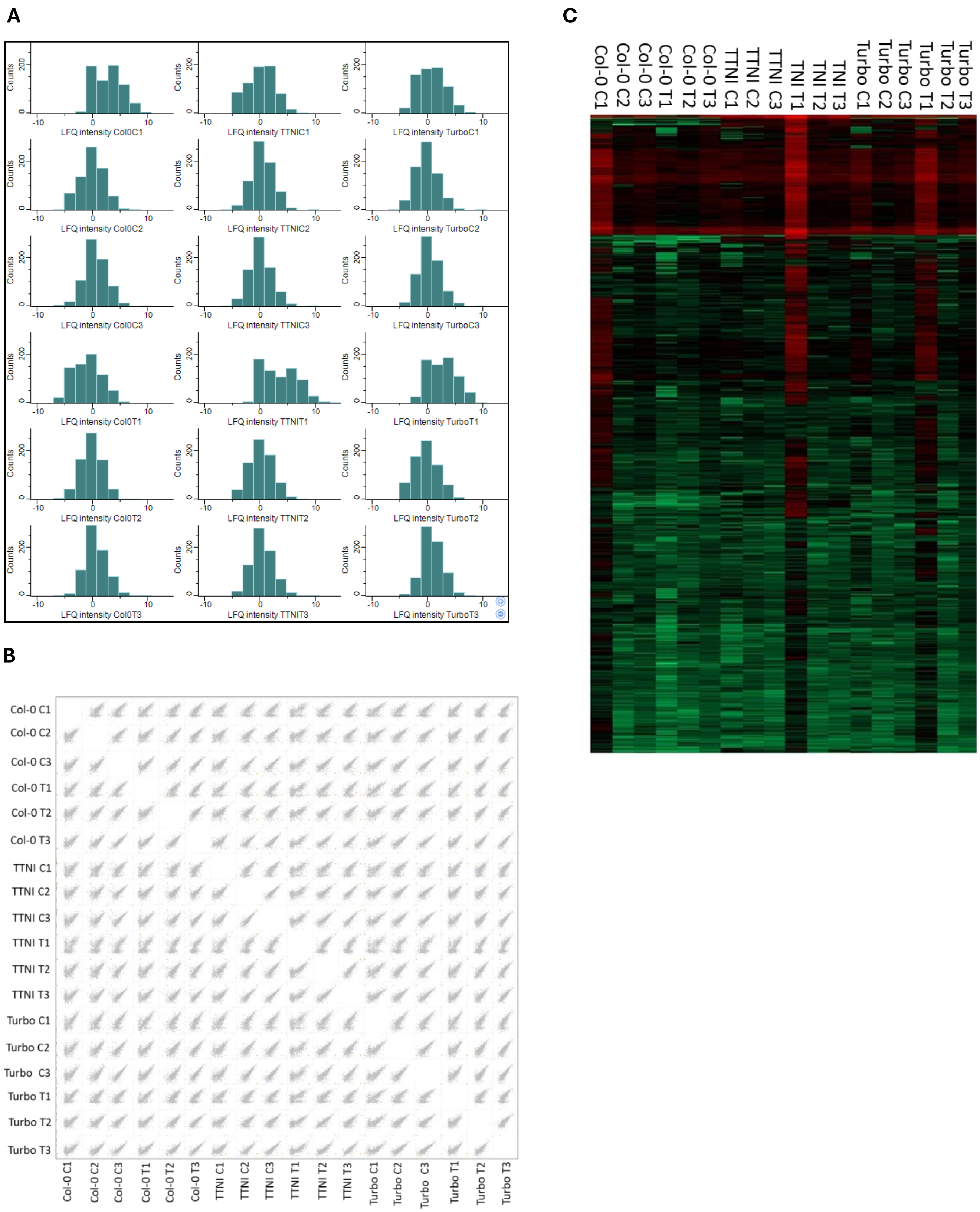


**Supp. Figure 4. Statistical analysis of the interacting proteins identified from AP-MS**. (A) Label-Free Quantification intensity plot and (B) multi-scatter plot showing relationship among all the samples and controls. Grey dots, interacting proteins; red dot, TNI. (C) Heat map of all the pulldown replicates. C stands for biotin untreated, T for biotin treated, TTNI for TbID-TNI.

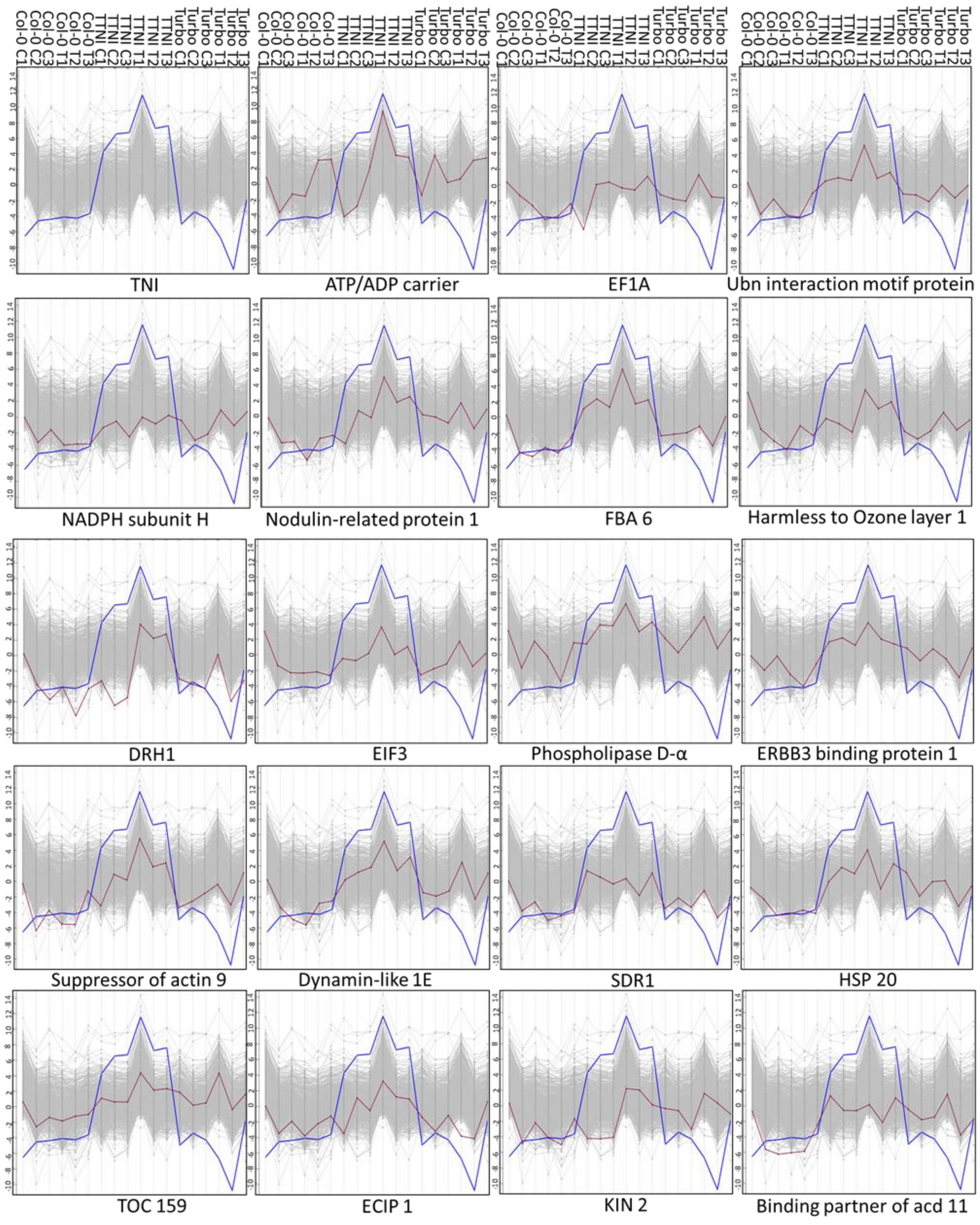


**Supp. Figure 5. Statistical analysis of AP-MS**. (A) LFQ intensity profile plot of interactors from the AP-MS. Blue line, TbID-TNI; red line, interactor labelled below each plot; grey lines, all the identified proteins in the AP-MS.C stands for biotin untreated, T for biotin treated, TNI for TbID-TNI.

A

| Sl.No. | TAIR ID | Proteins |
|---|---|---|
| 1 | AT1G43190 | Polypyrimidine tract-binding protein 3 |
| 2 | AT3G13750 | Beta galactosidase 1 |
| 3 | AT4G27640 | ARM repeat superfamily protein |
| 4 | AT5G49360 | Beta xylosidase 1 |
| 5 | AT5G56870 | Beta galactosidase 4 |
| 6 | AT2G36460 | Fructose-bisphosphate aldolase 6 |
| 7 | AT3G60190 | Dynamin-related protein 1E |
| 8 | AT4G24800 | EIN2 C-terminus interacting protein 1 |
| 9 | AT5G13490 | ADP/ATP carrier 2 |

B

| Sl.No. | TAIR ID | Proteins |
|---|---|---|
| 1 | AT1G43190 | Polypyrimidine tract-binding protein 3 |
| 2 | AT2G19830 | SNF7.2, VPS32-1 |
| 3 | AT2G41100 | Calmodulin4 |
| 4 | AT3G11730 | RAB GTPase Homolog D1 |
| 5 | AT4G08850 | MDIS1-interacting receptor like kinase2 (MIK2) |
| 6 | AT4G27640 | Importin-Beta 4 |
| 7 | AT4G39520 | Developmentally regulated g-protein (DRG1-1) |
| 8 | AT5G37780 | Calmodulin 1 |
| 9 | AT5G47520 | RAB GTPase homolog A5A |
| 10 | AT5G47230 | ERF5 |
| 11 | AT1G62300 | WRKY6 |
| 12 | AT4G37180 | UIF1 |
| 13 | AT1G18070 | EF1A |
| 14 | AT1G43690 | Ubiquitin interaction motif-containing protein |
| 15 | AT2G36460 | Fructose-bisphosphate aldolase 6 |
| 16 | AT3G01540 | DEAD box RNA helicase 1 |
| 17 | AT3G11400 | EIF3 |
| 18 | AT3G15730 | Phospholipase D alpha 1 |
| 19 | AT3G51800 | ERBB-3 Binding Protein 1 |
| 20 | AT3G59770 | SacI homology domain containing protein |
| 21 | AT4G02450 | HSP20-like chaperones superfamily protein |
| 22 | AT5G15970 | Stress responsive protein (COR6/KIN2) |
| 23 | AT5G16840 | Binding partner of acd11 1 |

**Supp. Figure 6. Overlap of TNI-interacting proteins identified in this study and the ubiquitinated proteins identified in the pulldown studies by (A) Kim et al., 2013 and (B) Song et al.., 2021.**
